# Ketamine Alters Tuning of Neural and Behavioral Spatial Working Memory Precision

**DOI:** 10.1101/2025.02.10.637233

**Authors:** Masih Rahmati, Flora Moujaes, Nina Purg Suljič, Jie Lisa Ji, Lucie Berkovitch, Kangjoo Lee, Clara Fonteneau, Charles H. Schleifer, Brendan D. Adkinson, Aleksandar Savič, Nicole Santamauro, Zailyn Tamayo, Caroline Diehl, Antonija Kolobaric, Morgan Flynn, Terry Camarro, Clayton E. Curtis, Grega Repovš, Sarah K. Fineberg, Peter T. Morgan, Katrin H. Preller, John H. Krystal, John D. Murray, Youngsun T. Cho, Alan Anticevic

## Abstract

Deficits in working memory (WM) are a hallmark of neuropsychiatric disorders such as schizophrenia, yet their neurobiological basis remains poorly understood. Glutamate *N*-methyl-D-aspartate receptors (NMDARs) are critical for spatial WM (sWM), with NMDAR antagonist ketamine known to attenuate task-evoked activation and reduce sWM accuracy. Cortical microcircuit models hypothesize that NMDAR antagonism impairs sWM by broadening neural spatial tuning, but this mechanism has not been directly tested in humans. Using a pharmacological fMRI approach, we showed how ketamine broadened neural spatial tuning, attenuated activation across visual, parietal, and frontal areas, and worsened sWM performance in healthy humans. Ketamine-induced changes in tuning were more consistent across individuals and brain regions than changes in overall activation and correlated with individual differences in sWM performance. These findings provide empirical evidence linking NMDAR antagonism to disruptions in cortical microcircuit dynamics, the resulting neural tuning alterations, and sWM impairments, advancing frameworks for therapeutic development.

## Introduction

Cortical computations—and consequently, key cognitive functions such as spatial working memory (sWM), the ability to temporarily maintain and manipulate spatial information—rely critically on N-methyl-D-aspartate receptors (NMDARs) in the brain. Disrupted NMDAR signaling has been implicated in mental illnesses such as schizophrenia (1, 2). Consistent with this, pharmacological disruption of NMDAR-mediated glutamatergic neurotransmission in healthy individuals induces symptoms resembling those of schizophrenia (3, 4). Specifically, the NMDAR antagonist ketamine has been shown to impair sWM—a hallmark cognitive deficit of schizophrenia—in both healthy humans (3) and non-human primates (5–7). Despite significant work done in this field, the precise mechanisms by which NMDAR antagonism disrupts sWM remain poorly understood.

Emerging evidence suggests that WM is supported by a distributed brain-wide network encompassing multiple hierarchical levels of neural organization, including cortical micro-circuits, local neural ensembles, and interconnected higher-order neural populations (8, 9). Electrophysiological recordings in monkeys have demonstrated persistent neural firing associated with sWM in the frontal cortex (5, 10). This persistent firing is thought to arise from recurrent excitatory connections within layer III microcircuits of the dorsolateral prefrontal cortex (dlPFC), predominantly mediated by NMDARs, and is further refined by inhibitory GABAergic interneurons to maintain spatial specificity (11, 12). Disruptions in this circuitry, particularly in NMDAR signaling, have been proposed as a key neural basis for sWM deficits (13, 14). Supporting this, studies have shown that the NMDAR antagonist ketamine disrupts the persistent neural firing of Delay cells in the monkey dlPFC, attenuating their activity and impairing sWM performance (5, 6).

Neurophysiologically-informed computational models have provided insights into sWM-related persistent representations in cortical microcircuits. These models suggest that interactions between glutamatergic lateral excitatory and GABAergic inhibitory connections modulate the fidelity of persistent neural representations (12, 15). Disruption of NMDAR signaling has been hypothesized to alter excitation-inhibition (E/I) balance, through perturbation of NMDARs on inhibitory interneurons, degrading the fidelity of neural representations, and impairing sWM performance (16).

Despite significantly advancing our understanding of how pharmacologically-induced NMDAR antagonism impacts neural processes, these theoretical and non-human primate findings offer limited insights. Specifically, they do not fully elucidate how microcircuit-level E/I imbalances disrupt the fidelity of sWM representations at the systems level and impair WM performance, particularly in humans.

Functional MRI (fMRI) in humans has identified sWM-related neural representations across sensory, association, and frontal cortices (17–21). Subanesthetic doses of ketamine in humans have been shown to reduce sWM task-related fMRI BOLD signals while increasing resting-state BOLD activity (4, 22, 23). Although these studies have advanced our understanding of the neural underpinnings of sWM and the deficits at the systems level, they provide limited information about how sWM representations are altered under NMDAR antagonism in localized structures, such as cortical columns. This limitation arises largely because current pharmacological neuroimaging approaches focus on overall activation within brain regions, neglecting the representational content of smaller, local neural ensembles (24– 26). Capturing this representational content is critical for bridging alterations in cortical microcircuits to systems-level changes and, ultimately, to the observed sWM deficits caused by NMDAR antagonism.

In this study, we address this limitation by investigating the effects of NMDAR antagonism on representational neural and behavioral measures of sWM in humans. More specifically, we empirically tested the hypothesis that NMDAR antagonism worsens sWM performance in healthy humans by broadening neural spatial tuning—the selective response profile of a neuron or ensemble of neurons to spatial stimuli. Using a pharmacological neuroimaging approach, we measured neural spatial tuning using fMRI BOLD signals collected during a sWM task under ketamine and placebo conditions. By employing Encoding Models (EM) (27, 28), we developed a framework for estimating voxel-level spatial tuning from sWM-related fMRI signals and quantifying changes induced by ketamine in healthy humans.

We found that ketamine broadens neural spatial tuning across spatially-selective brain regions and that this effect correlates with ketamine-induced deficits in sWM performance. These findings provide a mechanistic foundation for understanding how NMDAR antagonism impairs the fidelity of sWM representations in higher-order neural ensembles, leading to deficits in cognitive performance. Furthermore, ketamine attenuated sWM-related neural activation across the sWM network, consistent with previous findings in monkeys (5, 6) and humans (4). Interestingly, we found that compared to overall neural activation, ketamine-induced changes in neural tuning was more consistent across the brain. Collectively, this work introduces a novel integrative framework that bridges cortical microcircuit models, local neural ensembles, system-level neuroimaging readouts, and cognitive performance. This framework offers a robust foundation for developing therapeutic interventions targeting WM-related cognitive impairments.

## Results

Healthy participants (N=40) performed a spatial working memory (sWM) task, reporting the location of a memorized spatial cue after a delay. To examine NMDAR antagonist ketamine’s effects on sWM and neural activity, we collected fMRI BOLD signals during the task under both placebo and ketamine conditions (see methods and (26)). Using these BOLD signals, we identified spatially-selective brain regions, measured neural spatial tuning and activation, and compared them across conditions within these regions. Finally, we assessed the relationship between ketamine-induced neural tuning changes and sWM performance.

### Effect of ketamine on sWM performance

Through a within-subject design, we compared sWM performance, measured as response angular error in participants responses, under placebo and ketamine (Fig. 1). Angular error (θ_*AE*_) was decomposed into two components: i) sWM individual-level bias, defined as the average directional distance between cue and response locations, and ii) corrected angular error, defined as the angular distance between each trial’s cue and response locations subtracted by the bias. Compared to placebo, ketamine significantly increased angular error (0.89_*o*_, %95 CI: [0.62 Inf]), while it did not affect error variability across individuals—those with larger sWM errors under placebo also had larger errors under ketamine (Fig.1B, Pearson correlation *r* = 0.67, *p* < 0.001). Ketamine worsened sWM angular error in 31 out of the 40 participants (Fig. 1B, the ordered histogram plot). Additionally, ketamine increased bias on average (0.72_*o*_, %95 CI: [0.43 Inf]), while it did not affect the variability of bias across participants (Fig. S1). Moreover, we measured the consistency of each participant’s responses across repeated attempts through sWM dispersion, defined as the across-trials standard deviation of directional angular error. Ketamine significantly increased sWM dispersion (1.01_*o*_, %95 CI: [0.64 Inf]), while it did not affect dispersion variability across individuals (Fig. S1).

**Fig. 1.**
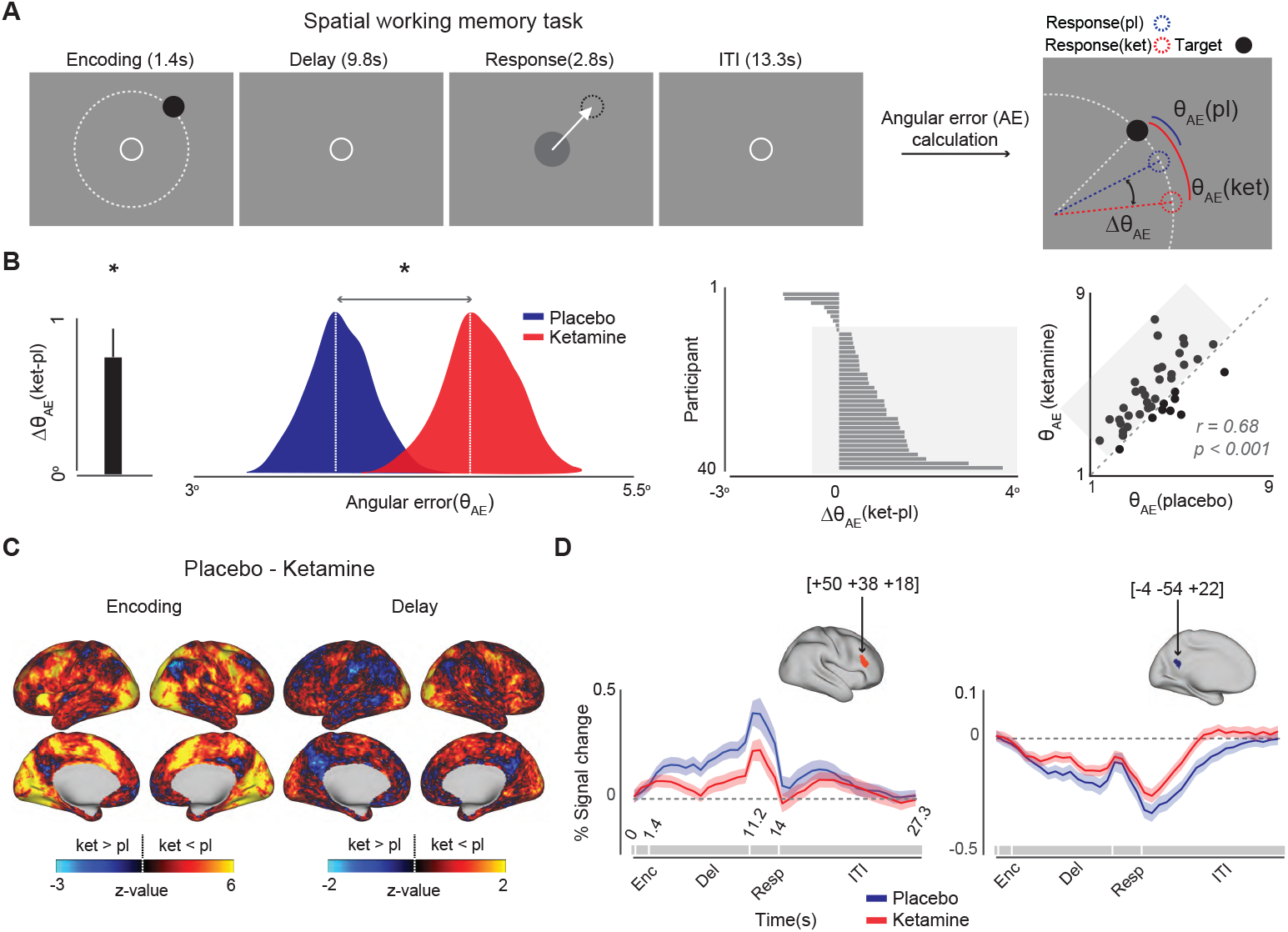
Spatial working memory task design, behavioral performance, and overall effect of ketamine on neural activation. **(A)** Spatial working memory (sWM) task; participants (N=40) reported the memorized location of a spatial cue (the black circle) after a delay period while keeping their fixation at the center of the display. At the end of the delay period, participants responded by positioning a joystick at the memorized location of the cue. sWM performance was calculated through the corrected angular error(*θ*_*AE*_), defined as the angular distance between subject’s response(the dashed circle) and the cue location(the solid circle). To remove the systematic angular bias that can arise on an individual level, we subtracted each participant’s trial-mean directional error from each trial’s uncorrected error. **(B)** Behavioral performance; The bar graph shows the group-mean ketamine-induced change in angular error, indicating that ketamine significantly increases the sWM error at the group level. The ridgeline plot shows the distribution of angular error across all participants under placebo and ketamine (dashed white lines indicate group means). The ordered histogram shows the worsening effect of ketamine on angular error across individual participants, indicating that 31 out of 40 participants had larger angular error under ketamine compared to placebo (the highlighted area). Each bar represents a single participant, and larger values indicate a stronger effect of ketamine on the angular error of sWM. The scatter plot depicts the relationship between angular error under ketamine and placebo conditions. Participants with larger angular error under placebo have larger error under ketamine, showing that ketamine does not affect the overall distribution of sWM performance across participants. The highlighted rectangle shows participants with larger angular error under ketamine, corresponding to the highlighted participants in the ordered histogram. **(C)** Ketamine-induced activation change; Ketamine-induced change in the activation measured as the difference between activation under placebo and ketamine (represented through the z-scored t-statistics). The warmer colors on the activation change map indicate stronger deactivating effects of ketamine relative to placebo. **(D)** Time course of neural activation; The time course of the group-level neural activation under placebo and ketamine within an area of interest (ROI) in the frontal (the left panel) and one in the precuneus cortices (the right panel). These exemplar ROIs were selected based on our previous findings on the effect of ketamine on neural activation (4) and functionally defined networks(29). Shaded areas in the plots correspond to one standard error across participants. Each ROI’s surface color is proportional to the overall ketamine-induced change in activation within the corresponding ROI.

### Ketamine weakens overall neural activation

Ketamine decreased neural activation across the brain. Using a general linear model (GLM), we calculated neural activation throughout the entire brain during the encoding and delay phases. We observed that ketamine weakens neural activity in most brain regions, except for the default mode network (DMN) (Fig. 1C). Additionally, we examined neural activation time series within selected functionally defined networks (29) (Fig. 1D). While ketamine attenuated neural activation in the dorsal attention and frontoparietal networks, it did not significantly affect DMN activity during the sWM task. These results robustly replicate previous findings on ketamine effects on neural activation (4). The group-mean activation in cortical and subcortical areas is shown in Fig. S2.

### Spatial selectivity and neural tuning can be reliably calculated from sWM task-evoked signals

We measured neural spatial tuning by estimating voxel-level receptive fields (RF) using an encoding model-based (EM) approach. An EM formulates the response of each voxel as a linear combination of a set of basis functions (BFs) each tuned to a specific visual field location (Fig. 2A). The regression weights in an EM determine the strength of the link between each voxel and each BF. Therefore, a voxel’s receptive field (RF) can be estimated as the linear sum of all BFs, each weighted by its corresponding regression weight (Fig. 2B). Since the visual field is a two-dimensional space, RFs, and BFs can be modeled as two-dimensional functions characterized by two parameters (i)the RF center (i.e., the visual field location they are tuned to), and (ii) the RF’s width, which is inversely proportional to the RF’s tuning (i.e., the wider the RF, the less tuned it is) (Fig. 2B). However, due to the fixed eccentricity of the stimuli used in this experiment, we simplified our computations without losing generality by using one-dimensional functions (of angular position) to model both the BFs and RFs (Fig. 2A and 2B).

**Fig. 2.**
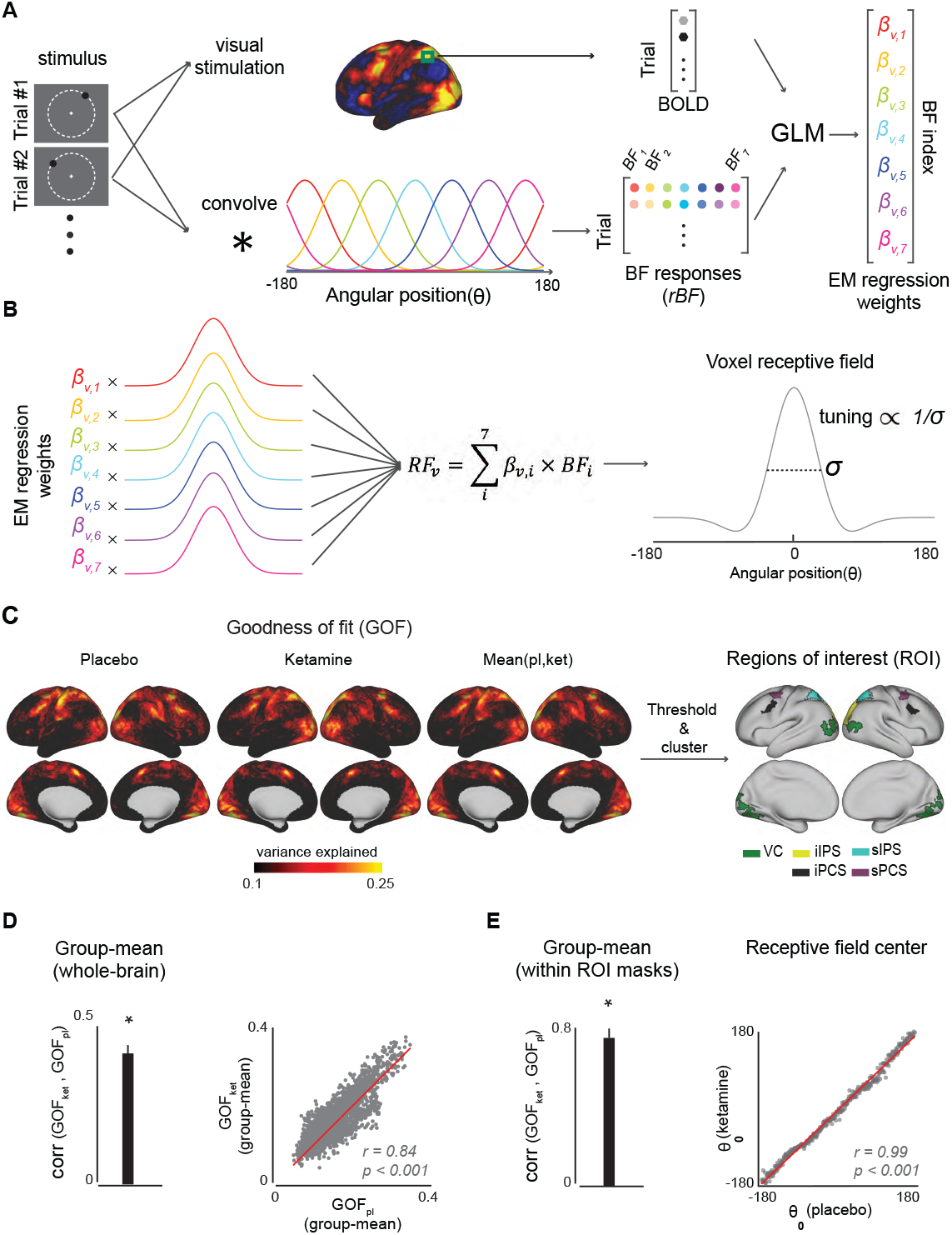
Encoding Model and tuning estimation. **(A)** Building the Encoding Model (EM); EM is a set of regression weights that determine the strength of the link between each voxel and a set of basis functions (BF set), selected based on the known tuning properties of the visual system. For simplicity, we represented the visual space as one-dimensional. This minimally affects the generality of our approach, due to the fixed eccentricity of the visual stimuli across trials. The regression weight between voxel *v* and the *i*-th BF (*β*_*v*,*i*_) is calculated through a General Linear Model (GLM) with two inputs: 1-the timeseries of the voxel’s response to a training set of visual stimuli (the upper path) and 2-the corresponding BF responses (rBF) to the cue location across trials in the training set (the lower path). **(B)** The Receptive Field (RF), or tuning profile, of each voxel is measured as the linear sum of all BFs, each weighted by the corresponding regression weight between the BF and that voxel (*β*_*v*,*i*_). Each RF is characterized through two parameters: i) the angular position of its peak (*θ*_0_), representing the angular position in the visual field that a voxel is most responsive to, and ii) its width (*σ*), which is inversely proportional to the tuning precision. **(C)** The spatial selectivity map and regions of interest (ROI); The spatial selectivity map consists of voxels that their response to a visual stimulus is sensitive to the location of the stimulus in the visual field. The sensitivity of voxels is measured as the goodness-of-fit (GOF) of the EM. ROIs are the most spatially-selective areas across the whole brain, determined as clusters of voxels which pass a GOF threshold applied to the condition-mean GOF map. Here we only focus on those clusters that are known for their spatial selectivity (30): visual cortex (VC), inferior intraparietal sulcus(iIPS), superior intraparietal sulcus (sIPS), inferior precentral sulcus (iPCS), and superior precentral sulcus (sPCS). **(D)** The consistency of spatial selectivity maps across conditions; The bar plot: group-mean Pearson correlation between GOF maps under placebo and ketamine, calculated independently for each participant. The error bar represents one standard error (SE) across participants. The scatter plot: the relationship between group-mean GOFs under placebo and ketamine and across all voxels. **(E)** The consistency of spatial selectivity across conditions within ROIs; The bar plot: group-mean Pearson correlation between GOFs under placebo and ketamine, calculated independently for each participant within a mask consisting of all ROIs combined. The error bar represents one standard error (SE) across participants. The scatter plot: The relationship between each voxels’ RF centers ((*θ*_0_) in panel B), estimated through two EMs built independently under placebo and ketamine. Each data point represents *θ*_0_ s of voxels with RF center within a 1^*o*^ range of angular position. Voxels are ordered according to their RF center under placebo. Voxels from all participants and within the combined ROIs mask are merged into one voxel pool. Notice that the highly close RF centers, estimated independently under two conditions, highlight the validity of this EM-based approach in estimating the spatial tuning.

We constructed two condition-specific EMs to estimate RFs from BOLD signals for all voxels across the brain under each condition. Additionally, we used these EMs to identify spatially-selective regions of interest (ROI), within which we compared tunings under placebo and ketamine. Identifying these ROIs is crucial because it allows us to validly assess ketamine’s effect on spatial tuning only across spatially-selective voxels. EMs are grounded in the spatial selectivity properties of voxels, meaning the quality of signal fitting depends on the amount of spatial information contained. Consequently, the goodness-of-fit (GOF) of an EM for each voxel can be interpreted as a measure of its spatial selectivity. GOF distributions across the entire brain were consistent between conditions and align with the distributions of spatial selectivity maps reported in previous studies (30, 31) (Fig. 2C). Notably, areas such as visual, intraparietal, and frontal cortices have higher GOF values compared to non-spatially selective areas (e.g., the default mode network and language areas). To select spatially-selective ROIs, we first averaged group-mean GOF maps across conditions and thresholded them to identify spatially-selective clusters across the whole brain. We then identified ROIs by selecting clusters that anatomically match previously reported areas involved in visual perception and WM (30, 32, 33). The distributions of GOF across participants and the relationship between across-participants GOF distributions under placebo and ketamine are demonstrated in Fig. S3A and S3B, respectively.

Importantly, the validity of between-condition comparison of tuning depends on the reliability and comparability of the RFs estimated under these conditions. We validated the reliability of the EM-estimated RFs in two ways. First, we tested the quality of EMs under both conditions at the individual and group levels. The bar plot in Fig. 2D shows significant individual-level correlation between GOFs maps under placebo and ketamine, averaged across participants (Pearson correlation *r* = 0.42, *p* < 0.001). Moreover, this correlation is significantly stronger within the spatially-selective ROIs (the bar plot in Fig. 2E) (Pearson correlation *r* = 0.78, *p* < 0.001). At the group-level, GOF maps are strongly correlated under placebo and ketamine (the scatter plot in Fig. 2D) (r = 0.84, *p* < 0.001). Finally, we found that the estimated angular positions to which each voxel is tuned (i.e., RF center θ_0_) are consistent across conditions (the scatter plot in Fig. 2E). These findings substantiate the validity of our subsequent tuning analysis conducted using these EMs.

### Spatially-selective regions can track trial-wise accuracy of sWM

A critical yet under-explored aspect of studying the neural mechanisms underlying WM and its deficits is identifying neural measures that can track trial-wise accuracy, thereby linking neural signals to cognitive performance. Two fundamental components of this approach are: (1) determining the appropriate neural measures, and (2) pinpointing the brain regions where these measures are indicative of cognitive performance. In this study, we tested the hypothesis that our EM-based spatial selectivity measure, specifically the EM’s goodness-of-fit (GOF), could identify regions of interest (ROIs) where neural activation, measured by the magnitude of the BOLD signal, can predict trial-wise sWM precision. We evaluated this hypothesis by comparing neural activation corresponding to subsets of trials ordered by their sWM angular error.

First, we divided each participant’s trials into ten deciles under each condition based on their sWM performance. As shown in Fig. 3A, sWM error is consistently higher under ketamine compared to placebo across all ten deciles. To maximize the contrast in sWM performance and test it against corresponding neural activation, we compared sWM error and neural activation for the placebo 1^st^ (the most precise trials across both conditions) and ketamine 10^th^ (the least precise trials across both conditions) deciles.

**Fig. 3.**
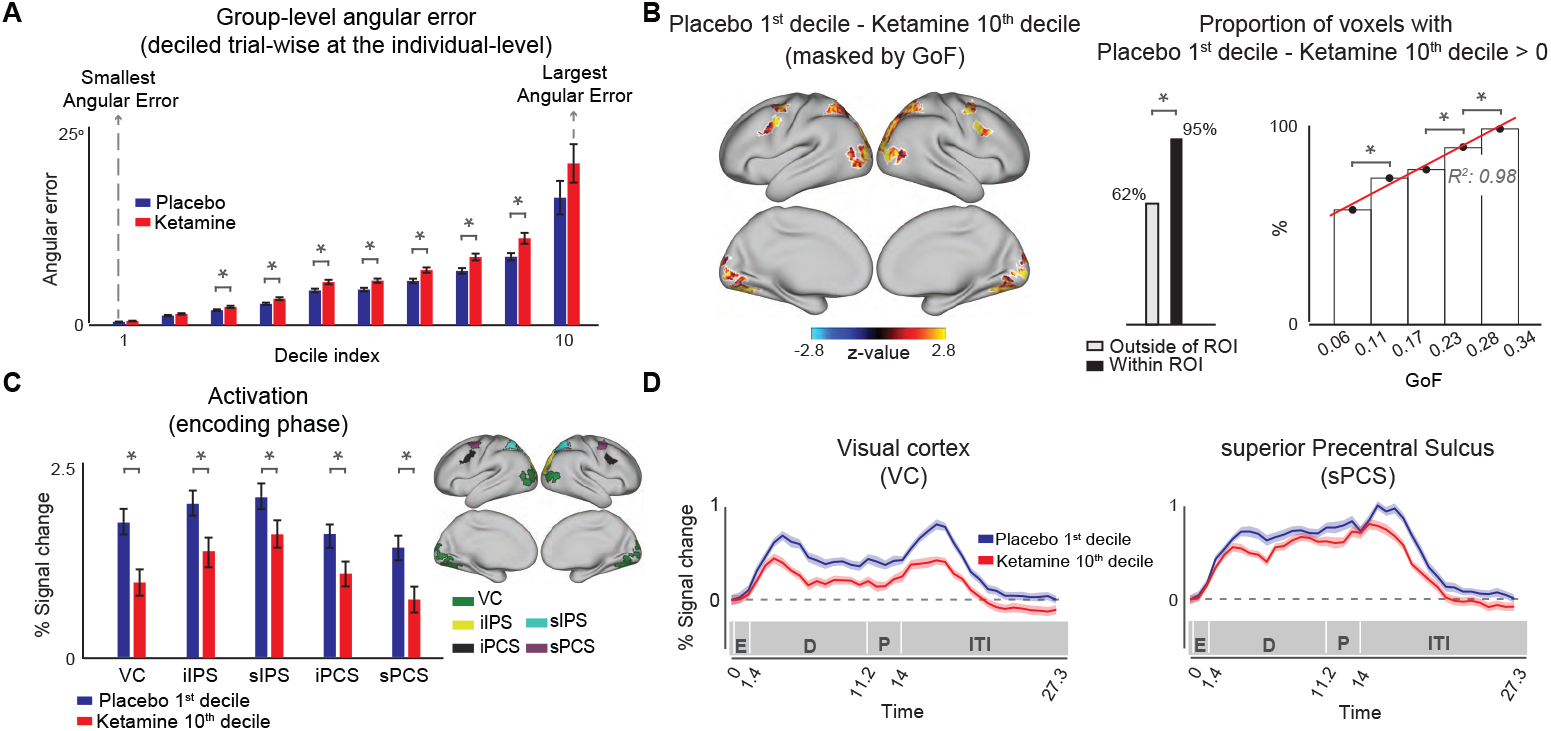
Association between the neural effects of ketamine and its effect on trial-wise sWM accuracy. We tested if neural activation can capture the variability in the trial-wise accuracy of sWM. **(A)** Trial-wise accuracy of sWM; For each participant, trials were divided into ten deciles according to their angular error. Blue and red bars depict the group-mean angular errors for different deciles under placebo and ketamine, respectively. Error bars show one standard error(SE) across participants. The asterisks show statistical significance. **(B)** The relationship between spatial selectivity and sWM accuracy-related activation; The brain surface depicts, within the GOF-based ROIs, the difference in neural activation between placebo 1^st^ and ketamine 10^th^ deciles, which corresponds to the largest trial-wise contrast in sWM accuracy. The left-side bar plot depicts the proportion of voxels with larger activation corresponding to placebo 1^st^ compared to ketamine 10^th^ decile. The right-side bar plot shows this proportion as a function of GOF. Each bar corresponds to the voxels, throughout the whole brain, within a specific GOF range. The height of each bar represents the percentage of voxels with larger activation of placebo 1^st^ compared to ketamine 10^th^ decile among voxels in the corresponding GOF range. **(C)** The group-mean activation for the placebo 1^st^(blue) and ketamine 10^th^(red) deciles within different ROI. Error bars show one standard error (SE) across participants. (D) The timeseries depict the temporal profile of the group-mean activation for the placebo 1^st^ (blue) and ketamine 10^th^ (red) over the course of a trial in one sensory (VC) and one frontal (sPCS) ROIs. Shaded areas show one standard error (SE) across participants.

To assess how activation contrast predicts maximum performance contrast within spatially-selective ROIs compared to the areas outside these ROIs, we compared the portion of voxels with positive activation contrast (i.e., placebo 1^st^ > ketamine 10^th^) within and outside ROIs. As shown in the middle bar plot in Fig. 3B, there is a 33% difference in the portion of voxels that respond more strongly during placebo 1^st^ trials, compared to ketamine 10^th^ trials (binomial test, p « 0.01). We also found that the portion of positive activation contrast linearly increases with GOF (R^2^ = 0.98) (Fig. 3B, right panel). These results suggest that there are spatially-specific mechanisms (i.e., mechanisms specific to spatially-selective areas) through which ketamine affects neural activation across the brain.

Moreover, we found that in all spatially-selective ROIs, neural activation is significantly larger for placebo 1^st^ trials, compared to ketamine 10^th^ trials (respectively for VC, iIPS, sIPS, iPCS, and sPCS: *Mean*_*pl*_ - *Mean*_*ket*_ = [0.77, 0.62, 048, 0.51, 0.66], SE = [0.19, 0.22, 0.20, 0.17, 0.18], p = [0.001, 0.004, 0.01, 0.002, 0.0004]) (Fig. 3C). Fig. 3D illustrates the time series of the BOLD signal within the visual cortex (VC) and superior precentral sulcus (sPCS), highlighting the difference in neural activation between the placebo 1^st^ and ketamine 10^th^ deciles throughout the course of a trial. These findings underscore the potential of EM-based spatial selectivity measures in identifying brain regions where neural activation is predictive of cognitive performance on a trial-wise basis. Neural activation corresponding to different deciles and conditions are shown in Fig. S4.

### Ketamine broadens spatial tuning in multiple brain areas

Our main hypothesis in this study was that the NMDAR antagonist ketamine decreases neural spatial tuning, reflected in broader RFs under ketamine compared to placebo (Fig. 4A). To test this, each voxel’s RF width was calculated as the inverse of the concentration parameter in a von Mises function, to which the RF was fitted (Fig. 2B). We independently calculated RF width under each condition for all voxels within the spatially-selective ROIs. The middle column brain map in Fig. 4B shows the group-mean ketamine-induced change in RF width, averaged across all voxels within each ROI. The left side bar plot in Fig. 4C quantitatively illustrates the ketamine-induced change in RF width for all participants, where ketamine significantly broadened spatial tuning in the visual cortex (VC) (*σ*_*ket*_ - *σ*_*pl*_ = 3.33°, *p* < 0.003) and the inferior intraparietal sulcus (iIPS) (*σ*_*ket*_ - *σ*_*pl*_ = 4.41°, *p* < 0.01). Additionally, ketamine broadened tuning in suprior precentral sucluc (sPCS) almost significantly (*σ*_*ket*_ - *σ*_*pl*_ = 3.57°, p = 0.06). These results support our hypothesis that ketamine decreases neural spatial tuning by broadening RFs in specific brain regions. The distribution ketamine-induced change in spatial tuning across participants and ROIs are shown in Fig. S5.

**Fig. 4.**
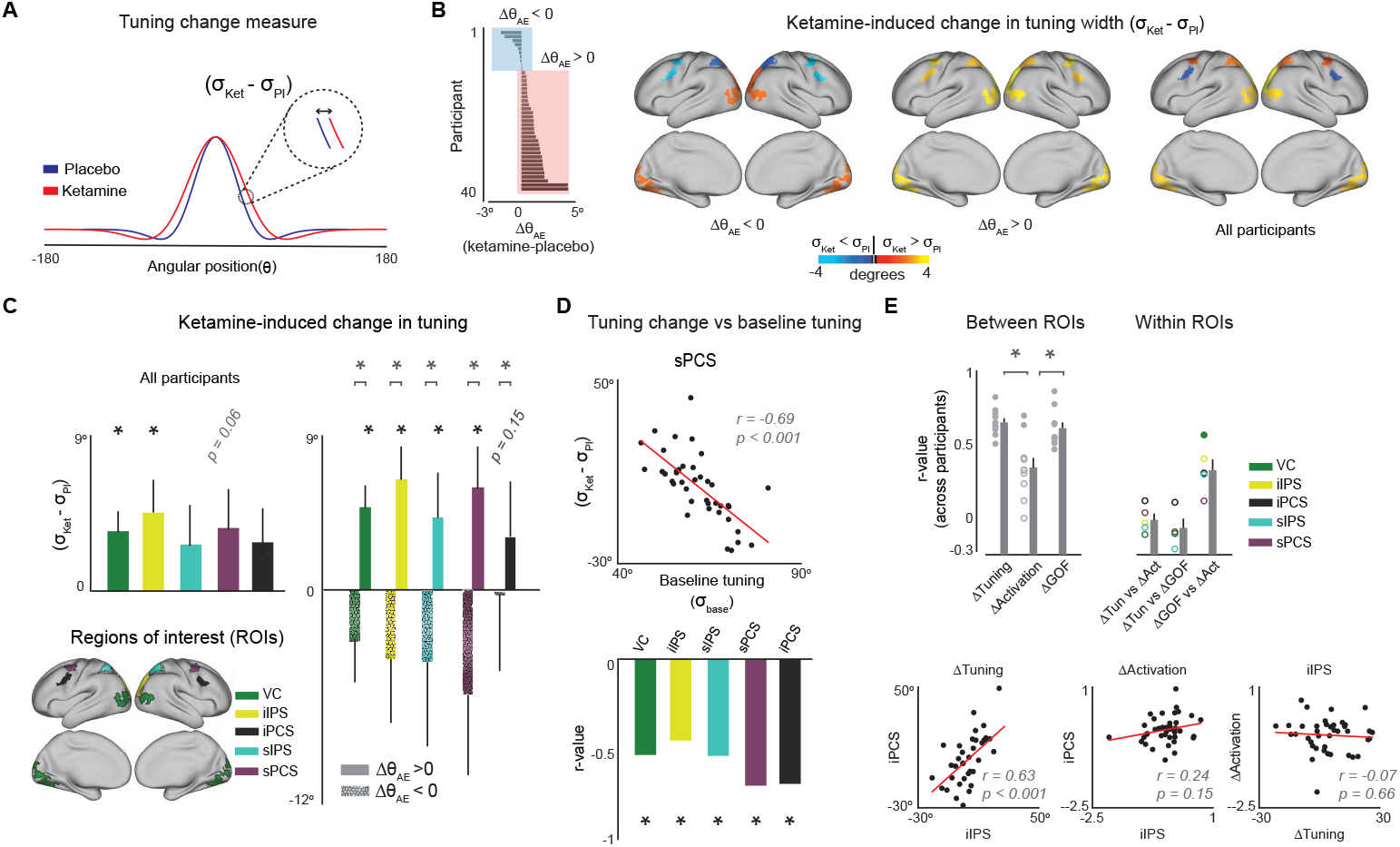
Effect of ketamine on neural tuning. **(A)** Schematic showing the tuning change measure; The effect of ketamine on tuning is measured as the difference in the width of the EM-estimated voxel’s receptive field (RF) (*σ*), under placebo and ketamine. **(B)** Ketamine effect on tuning; The group-average effect of ketamine on tuning for three subsets of participants according to the effect of ketamine on their sWM Angular Error (AE): i) Δ*θ*_*AE*_ < 0: participants who had less precise sWM performance under placebo, ii) Δ*θ*_*AE*_ > 0: participants who had less precise sWM performance under ketamine, and iii) all participants. The brain surfaces show the group-mean effect of ketamine on tuning averaged across all voxels within each ROI. **(C)** Quantitative comparison of ketamine effect on tuning; Across all participants, ketamine significantly broadened tuning in VC and iIPS, and almost significantly in sPCS. Moreover, among those participant with worsening ketamine effect on their sWM (i.e., Δ*θ*_*AE*_ > 0), ketamine significantly broadened tuning in VC, iIPS, and sPCS, and almost significantly in sIPS. However, ketamine did not affect tuning among participants with Δ*θ*_*AE*_ < 0. **(D)** The relationship between ketamine-induced change in spatial tuning and baseline tuning (i.e., tuning under placebo); The scatter plot depicts the tuning change as a function of baseline tuning in sPCS, as an example ROI. Each data point represents one participant. The bar plot represents the correlation between tuning change and baseline tuning within different ROIs, showing that ketamine had a smaller broadening effect in participants with larger baseline tuning. **(E)** The between-ROIs relationship between ketamine-induced changes in tuning, the magnitude of activation, and spatial selectivity (GOF), as well as the within-ROI relationship between these measures; The left set of barplots show the across-participants Pearson correlation between all possible pairs of ROIs for ketamine-induced changes in tuning, neural activation, and EM’s GOF. Each measure is calculated as its average across all voxels in each ROI. Each gray dot corresponds to one of the ten possible pairs of ROIs. Solid and empty dots show significant and insignificant correlations, respectively (corrected for multiple comparisons). The right set of barplots represent the Pearson correlations between ketamine-induced changes in tuning, neural activation, and GOF within each of the five ROIs. The scatter plots depict the between-ROIs relationships for two example ROIs for ketamine-induced changes in tuning and neural activation, as well as the within-ROI relationship between these two changes in an example ROI. Each dot in the scatter plots corresponds to one participant.

### Alterations in neural tuning predicts sWM performance

Biophysically-inspired computational models of cortical microcircuits hypothesize that the NMDAR antagonism worsens sWM performance by broadening neural spatial tuning. To test the relationship between ketamine’s impact on spatial tuning and sWM accuracy, we measured the broadening effect of ketamine on tuning in two subgroups of participants based on their ketamine-induced change in sWM precision (i)the 31 participants with worsened sWM under ketamine, and (ii) the 9 participants with decreased sWM angular error (see the ordered histogram plot in Fig. 4B). We found that among participants with weaker sWM performance, spatial tuning was significantly broader under ketamine in all ROIs except iPCS; for VC, iIPS, sIPS, and sPCS, respectively: *σ*_*ket*_ - *σ*_*pl*_ = [4.93°, *p* < 0.001], [6.49°, *p* < 0.005], [4.28°, *p* < 0.05], [6.02°, *p* < 0.02]. Additionally, ketamine broadened tuning in iPCS (3.23°, p = 0.15). However, ketamine did not significantly affect spatial tuning in any of the ROIs among participants with worse sWM under placebo, compared to ketamine (Fig. 4C, the right error bar plot).

Moreover, we tested the relationship between the ketamine-induced changes in spatial tuning and each participant’s baseline tuning (i.e., tuning under placebo). We found that the broadening effect of ketamine on spatial tuning is smaller in participants with larger baseline tuning (Fig. 4D).

### Ketamine affects neural activation and spatial tuning differently

As shown in Fig. 3C and Fig. 4C, ketamine decreases both neural activation and spatial tuning in spatially-selective ROIs. An important question, then, is if these two effects arise from a shared cortical mechanism. To answer this question, we first tested the consistency of ketamine-induced changes in neural tuning and activation across the brain. Specifically, we calculated the across-participants Pearson correlations between ketamine-induced changes in each of these measures across different ROIs. As depicted in the upper left panel of Fig. 4E, ketamine-induced changes in neural tuning were highly correlated across ROIs, with Pearson correlation coefficients (r) ranging from 0.65 to 0.75 (p < 0.05, corrected for multiple comparisons), suggesting a brain-wide effect of ketamine on neural tuning. In contrast, correlations for ketamine-induced changes in neural activation during the encoding phase of the sWM task were not significant across ROIs, except between iIPS and VC, as well as between iIPS and sIPS. This suggests that, unlike tuning, this effect was not consistent across the brain (the upper left panel in Fig. 4E). Cross-correlations across ROIs are shown in Fig. S5C.

Finally, we examined the across-participants relationship between ketamine-induced changes in neural tuning and overall activation, neural tuning and GOF, and GOF and overall activation within each spatially-selective ROI. We found that ketamine-induced changes in tuning were not correlated with changes neither in neural activation, nor in spatial selectivity, measured as GOF, in any of the ROIs (the upper right panel in Fig. 4E). This reinforces the reliability of our measures, showing that ketamine-induced changes in tuning are not substantially influenced by variability in the identification of spatially-selective voxels. Furthermore, consistent with these across-participants correlations, we did not observe any voxel-wise relationship between ketamine-induced changes in neural activation and spatial tuning within individual ROIs (Fig. S5D; Fig. S6B). These findings suggest that distinct cortical microcircuit mechanisms may underlie the effects of ketamine on attenuating resting-state neural activation and broadening spatial tuning.

## Discussion

The overarching goal of this study was to test the hypothesis that sWM deficits induced by NMDAR antagonism can be explained mechanistically through ketamine’s broadening effect on spatial tuning. Our findings revealed that the administration of ketamine, a NMDAR antagonist, is associated with both behavioral and neural signatures that support this prediction. Our findings replicate prior work showing that sWM performance deteriorates under the influence of ketamine (4, 5). This behavioral impairment dovetailed with our neuroimaging results, which show a reduction in spatial tuning across multiple spatially-selective brain areas, including regions of the visual, parietal, and prefrontal cortices. Moreover, ketamine-induced alterations in neural tuning predicted the changes in sWM performance due to ketamine administration. Finally, we found that the magnitude of neural activation in spatially-selective regions can capture the individual-level, trial-wise variations in sWM precision. These results provide the first empirical evidence for the hypothesis developed by biophysically-grounded computational models of cortical microcircuit proposing tuning alteration as a mechanism underlying sWM deficits under NMDAR antagonism (16). Bridging the gap between the effects of NMDAR antagonism on cortical microcircuitry and system-level changes—ultimately leading to sWM impairments—is crucial for developing comprehensive pharmacological models of mental illnesses like schizophrenia, providing deeper insights into their neural basis, and guiding the creation of novel diagnostic tools and therapeutic interventions.

From a mechanistic standpoint, NMDARs are integral to synaptic plasticity and play a pivotal role in modulating cortical microcircuit dynamics (2, 34, 35). Our hypothesis regarding decreased neural tuning under NMDAR antagonism is inspired by two key lines of evidence: i) Electrophysiological recordings showing that NMDAR antagonism decreases sWM-related persistent neural firing and tuning indices in the dlPFC of non-human primates (5, 6), and ii) Biologically inspired computational models of cortical microcircuits, which hypothesize mechanistic explanations for how NMDAR antagonism could broaden neural tuning (16). Our findings complement those in non-human primates and provide empirical support for predictions made by cortical microcircuit models. Specifically, we observed both attenuated sWM-related activity and broadened neural spatial tuning due to NMDAR antagonism in humans. However, critical differences exist between our findings, those observed in non-human primates, and the predictions of computational models, warranting further investigation.

### Relation to ketamine-induced changes at the cellular level

The effects of NMDAR antagonism on neural firing and tuning indices observed in monkeys (5–7) were measured from a subset of dlPFC neurons, limiting insights into the dynamics of larger neural ensembles. For instance, in (5), neural firing was recorded exclusively from pyramidal Delay and Response cells in dlPFC layers III and V, respectively, which exhibit opposite responses to ketamine administration (i.e., decreased firing in Delay cells versus increased activity in Response cells). However, due to the small number of neurons sampled in these studies, it is challenging to derive a holistic measure representing the aggregate effects of NMDAR antagonism across various cell types on the tuning of local neural ensembles. In contrast, our findings address this critical gap by measuring ketamine-induced changes in neural tuning at the voxel level, capturing the activity of cortical circuits more comprehensively. This approach provides a broader and more integrative perspective on the impact of NMDAR antagonism on neural tuning in humans.

### Relation to the predictions of ketamine-induced effects by computational models of cortical microcircuits

Biologically-inspired computational models of cortical microcircuits predict that NMDAR antagonism weakens lateral inhibition from inhibitory interneurons to pyramidal cells, leading to broader neural representations of sWM and, ultimately, degraded sWM performance (16). While our findings align with this mechanistic explanation, a critical distinction exists between our results and these theoretical models. These cortical microcircuit models are based on a network architecture spanning the entire stimulus space, where neurons represent all possible stimulus locations (15, 16). Consequently, the simulated neural tuning reflects the population representation of the entire stimulus space, analogous to neural population representations in retinotopically-organized cortical regions (20, 21). In contrast, our study, for the first time, tested the effects of NMDAR antagonism on neural tuning within local neural ensembles (i.e., voxels), each tuned to a specific spatial location. These local ensembles serve as the foundational building blocks of the broader neural populations simulated by theoretical models. This distinction is crucial because our findings provide strong empirical support for a more comprehensive mechanistic framework, bridging the effects of NMDAR antagonism across different levels: cellular neural firing, tuning within local ensembles, and systems-level neural populations. Ultimately, this multi-level perspective enhances our understanding of how NMDAR antagonism impacts neural representations of sWM and its relationship to sWM deficits.

### Implications for neural analysis

Many cognitive functions, such as sWM, rely heavily on representational neural measures—those that encode specific information about different stimulus values, such as neural tuning or population representations. In contrast, non-representational neural measures, like overall BOLD signal, reflect general neural activations without encoding specific stimulus-related information. While non-representational measures provide insights into overall brain activity or task engagement, representational measures are more effective for investigating the neural mechanisms underlying cognitive functions and their impairments. For instance, previous neuroimaging studies, primarily focused on overall neural activation, have shown opposite effects of NMDAR antagonism neural activation during resting-state (increased BOLD) and sWM task (decreased BOLD) (22–24, 36, 37). In contrast, our findings advance our understanding of neural disruptions caused by NMDAR antagonism through employing neural tuning as a more directly relevant measure.

Encoding Models (EMs) have recently led to significant advancements in understanding the neural mechanisms underlying various cognitive functions (38–40). By reconstructing neural population representations from neuroimaging signals, these biologically-informed models have demonstrated that sWM performance depends on the fidelity of higher-order memory representations across different neural populations within the WM network (32). In this study, we highlight the potential of EM-based approaches to elucidate the mechanisms underlying cognitive impairments. Specifically, we leveraged an often-overlooked capability of EMs: measuring neural tuning at the single-voxel level. This method is particularly valuable for investigating the complex neural disruptions associated with neuropsychiatric disorders. For instance, in light of our EM-based findings, we can speculate that the ketamine-induced broadening of spatial tuning within spatially selective cortical areas may not only impair the encoding of sWM but also affect a broader range of cognitive functions that rely on the integrity of these microcircuits.

### Relationship between neural tuning and neural activation

Another important aspect of our findings is that neural tuning may provide more information about inter-individual differences compared to overall neural activation. The responses of stimulus-selective neurons or voxels are influenced by various factors, including the strength of sensory input (e.g., the contrast of a visual cue), long-range task-dependent interactions (such as top-down attentional signals), and their intrinsic tuning properties. However, neural tuning is primarily a physiological property governed by cortical micro-circuitry. Therefore, it serves as a more reliable measure for investigating individual differences in response to brain-wide modulations, such as those induced by NMDAR antagonism. Our findings support this hypothesis. Ketamine-induced changes in neural tuning are highly correlated across different ROIs, suggesting that the inter-individual relationship regarding ketamine-induced changes in neural tuning is consistent across the brain. In contrast, we did not observe such correlations for overall neural activation within the same ROIs, underscoring the limitations of overall activation as a measure for capturing across-individual differences in response to NMDAR antagonism. This limitation becomes even more evident when considering that our EM-based approach provided a data-driven method to identify spatially selective voxels—and thus ROIs—independently for each participant using signals from the sWM task. This approach yields more precise subject-specific task-related measures, including those not directly linked to tuning, thereby substantially improving the reliability of our findings.

### Relationship between neural tuning and impairments in sWM

The ultimate goal of studying the neural basis of cognitive deficits is to develop comprehensive mechanistic explanations that link neural dynamics to cognitive impairments. NMDAR hypofunction has been implicated as a potential mechanism underlying cognitive deficits in schizophrenia, a prototypical psychosis spectrum disorder (1, 2, 41, 42). The broadening of spatial tuning, induced by NMDAR antagonist ketamine, and the associated impairment in sWM performance in our study mirrors the cognitive disturbances seen in psychosis spectrum disorder. This offers a model for further investigation towards developing new targeted cognitive remediation therapies. For instance, by using fMRI and spatial tuning metrics as biomarkers, clinicians might be able to identify individuals at high risk for psychosis who exhibit subtle changes in neural processing before full-blown symptoms emerge, allowing for earlier intervention.

Our findings provide empirical support for mechanistic models connecting cortical microcircuitry to the function of local neural ensembles, and sWM performance. However, achieving a more comprehensive explanation of how NMDAR antagonism leads to sWM deficits requires extending these investigations to examine these effects on sWM representations in larger neural populations and their relationship to cognitive performance. Recent EM-based analytics have successfully linked neural population representations to sWM performance in healthy humans under normal conditions (32). Therefore, a critical future direction is to examine how alterations in the tuning properties of local neural ensembles impact the fidelity of sWM representations at the population level and how these changes ultimately contribute to sWM deficits.

In conclusion, these findings highlight the potential of using pharmacological models combined with advanced neuroimaging techniques to simulate and study the neural underpinnings of sWM deficits in neuropsychiatric disorders. Such integrative approaches are crucial for moving beyond symptom-based characterizations of cognitive dysfunction towards a deeper understanding of the neurobiological processes that contribute to these deficits, ultimately informing more effective interventions and preventative measures for neuropsychiatric disorders.

## ACKNOWLEDGEMENTS

…

## Methods

### Participants

Forty healthy participants were recruited from the New Haven area via flyers and online ads and provided informed consent approved by Yale University Institutional Review Board. The eligibility of participants to receive ketamine was determined by the following criteria, as determined by a detailed telephone interview and an in-person clinical assessment: i) Age 21-60; ii) IQ>70 as measured via Wide Range Achievement Test (WRAT-3) and the Wechsler Adult Intelligence Scale (WAIS-III); iii) intact or corrected-to-normal vision; iv) weight < 300 lbs.; v) MR safe (free of metallic objects and absence of claustrophobia); vi) no serious medical or physical conditions, as confirmed by a selfreport, electrocardiogram, blood work, and physical examination by a licensed physician; vii) no lifetime neurological or psychiatric diagnoses; viii) minimal alcohol intake and no use of psychoactive drugs or history of abuse/dependence, confirmed by interview and urinalysis; ix) no first-degree relatives with DSM Axis-I diagnoses or alcohol/substance abuse history; x) no known sensitivity to ketamine or heparin; xi) no donation of blood in excess of 500ml within 2 months of participation.

### Task Design

Each participant completed a spatial Working Memory (sWM) task under two conditions; administration of ketamine and placebo (saline). In each trial of the sWM task, a cue was first presented for 1.4 seconds at a pseudorandomized location around an imaginary circle (fixed radius = 415 pixels) and participants were instructed to maintain its position in their mind over time (Fig. 1A). After a 9.8 seconds delay, a probe signaled participants to move the fMRI-compatible joystick to where they best remembered the cue location. The main behavioral measure, angular error (AE), was the angular distance between the initial cue placement and participants’ probe placement (Fig. 1A). We specifically used angular error due to the fixed eccentricity of cue locations. Before the sWM task, each participant completed a delayed motor-only task identical to the sWM task except that there was no sWM element (i.e., there was no instruction for maintaining and reporting the memorized cue location and participants moved the joystick to the cued location, which was re-cued during the response period). Each fMRI scan consisted of ten trials and each participant completed one and six blocks of motor and sWM tasks, respectively, under each of the placebo and ketamine conditions (within-subject design), resulting in 70 trials total in each condition. The task was presented on a Dell laptop with an LCD screen (1280 × 1024 pixel resolution) using E-Prime 2.0 software and analyzed using Matlab (R2023a).

### Ketamine study design

The study used a single-blind, within-subjects design, and all participants were blinded to treatment order. During the first scan session, saline placebo was administered, and in the second scan session, ketamine (initial bolus 0.23 mg/kg, continuous infusion 0.58 mg/kg/hr) was administered. Participants were not informed of the order of administration, though all were able to correctly identify ketamine administration. Counterbalancing was not done due to residual effects of ketamine administration. For further details on ketamine study infusion, please see (26).

### Behavioral performance

We used the angular distance between the initial cue presentation and probe placement as the primary measure for each participant’s sWM performance. The selection of *angular* distance was based on the fixed eccentricity of the cue locations used in the task, and simplified neural analysis without losing generality (particularly in building the encoding model and receptive field analysis); using Euclidean distance did not significantly affect any of the results. Furthermore, we decomposed angular distance into two components: (i) systematic angular bias, defined as the trial-averaged angular distance between the actual visual cue and the participant’s response, and (ii) trial-specific sWM angular error (AE), quantified as the angular distance between the memorized location and the actual cue location, accounting for the systematic bias (i.e., after the systematic bias is subtracted). Angular error is inversely proportional to sWM *accuracy*, showing how accurate a subject’s sWM is at each trial. We calculated the overall individual-level angular error as the absolute AE, averaged across all trials after excluding outlier trials. The exclusion criteria and other details of the behavioral measures follow the ones in (43). Additionally, we measured the *precision* of each participant’s sWM through response dispersion, calculated as the across-trials standard deviation of directional angular errors. Dispersion is inversely proportional to the individual-level overall precision and reflects the consistency of each participant’s sWM across repeated attempts.

### fMRI data acquisition

All data were obtained from a 3 Tesla Siemens scanner at the Yale Magnetic Resonance Research Center in New Haven, CT using a 32 (n = 13) or 64 (n = 20) channel phased array head coil. Prior to acquiring BOLD images, high-resolution T1w and T2w structural images were collected with 0.8 mm isotropic voxels in 224 ACPC aligned slices, in compliance with the adult Human Connectome Project (HCP) acquisition protocol. T1w images were collected with a magnetization-prepared rapid gradient-echo (MP-RAGE) pulse sequence (TR = 2400 ms, TE = 2.07 ms, flip angle = 8°, field of view = 256 × 256 mm). T2w images were collected with SPC (SPACE: sampling perfection with application-optimized contrasts using different flip angle evolution (Siemens)) (pulse sequence [TR = 3200 ms, TE = 564 ms, flip angle mode = T2 var, field of view = 256 ×256 mm]. A pair of reverse phase-encoded spin-echo field maps (anterior to posterior, and posterior to anterior) was also collected to aid distortion correction in preprocessing (voxel size = 2.5 mm isotropic, TR = 7220 ms, TE = 73 ms, flip angle = 90°, field of view = 210 ×210 mm, bandwidth = 2290 Hz). All BOLD images were acquired with specifications based on the adult HCP acquisition protocols at the time of collection. 54 interleaved axial slices were collected parallel to the anterior-posterior commissure (AC-PC) with 2.5 mm isotropic voxels using a multi-band accelerated fast gradient-echo, echo-planar sequence (acceleration factor = 6, time repetition (TR) = 700 ms, time echo (TE) = 31.0 ms, flip angle = 55°, field of view = 210 × 210 mm, matrix = 84 ×84, bandwidth = 2290 Hz). Each run was 9.33 min with 800 vol. Single-band reference images acquired before each BOLD run aided registration during preprocessing.

### fMRI data processing

Neuroimaging data was preprocessed using the Human Connectome Project (HCP) minimal preprocessing pipeline (44). A summary of the HCP pipelines is as follows. First T1/2-weighted structural images were corrected for bias-field distortions and then warped to the standard Montreal Neurological Institute-152 (MNI-152) brain template in a single step, through a combination of linear and non-linear transformations via the FMRIB Software Library (FSL) linear image registration tool (FLIRT) and non-linear image registration tool (FNIRT) (45). Next, FreeSurfer’s recon-all pipeline was used to segment brain-wide gray and white matter to produce individual cortical and subcortical anatomical segmentations (46). Cortical surface models were generated for pial and white matter boundaries as well as segmentation masks for each subcortical grey matter voxel. Using the pial and white matter surface boundaries, a ‘cortical ribbon’ was defined along with corresponding sub-cortical voxels, which were combined to generate the neural file in the Connectivity Informatics Technology Initiative (CIFTI) volume/surface ‘grayordinate’ space for each individual participant (44). BOLD data were motion-corrected by aligning to the middle frame of every run via FLIRT in the initial NIFTI volume space. In turn, a brain mask was applied to exclude signal from non-brain tissue. Next, cortical BOLD data were converted to the CIFTI gray matter matrix by sampling from the anatomically-defined gray matter cortical ribbon and subsequently aligned to the HCP atlas using surface-based nonlinear deformation (44). Subcortical voxels were aligned to the MNI-152 atlas using whole-brain non-linear registration and then the Freesurfer-defined subcortical segmentation applied to isolate the subcortical grayordinate portion of the CIFTI space.

To reduce the potential artifactual, movement-driven change in the signal, we performed movement scrubbing (47) after the HCP minimal preprocessing pipelines. This can be done by i) computing statistics that reflect movement and its artifactual properties and flagging the problematic frames, and ii) ignoring or interpolating the problematic frames in analyses. As in prior work (4), all BOLD image frames with possible movement-induced artifactual fluctuations in intensity were flagged using two criteria: frame displacement (the sum of the displacement across all six rigid body movement correction parameters) exceeding 0.5 mm (assuming 50 mm cortical sphere radius) and/or the normalized root mean square (RMS) (calculated as the RMS of differences in intensity between the current and preceding frame, computed across all voxels and divided by the mean intensity) exceeding 1.6 times the median across scans. Any frame that met one or both of these criteria, was replaced with spline interpolated values based on neighboring good frames. Next, a high-pass filter (threshold 0.008 Hz) was applied to the BOLD data to remove low frequency signals due to scanner drifts. Finally, functional images were smoothed with a 2 mm FWHM Gaussian kernel.

### fMRI analysis

General linear models (GLM) were calculated for each participant using in-house software (Qu|Nex, https://qunex.yale.edu) implemented in Matlab (R2023a), with 12 motion regressors of no interest, baseline and drift regressors for each run. For assumed response modeling, task regressors for each epoch (i.e., encoding, delay, and probe) and condition were included. The hemodynamic response function (HRF) was modeled using an assumed boxcar function convolved with a gamma function (48). The Encoding Model (EM) GLM was calculated simultaneously with the task events GLM.

### Encoding Model and spatial tuning

We measured the neural spatial tuning by estimating the Receptive Field (RF) of each voxel through Encoding Models (EM) (27, 28). EM estimates a voxel’s RF based on two assumptions. First, that the activity of each neuron is a linear combination of the outputs of a set of basis functions (BF), or information channels, each tuned to a specific spatial location in the visual field. Second, that the activity of each voxel is a weighted sum of the activity of all neurons in that voxel. As a result, the activity of each voxel is a linear combination of the response of the basis functions to a given spatial location. We can formulate this mathematically as:

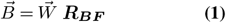

Where:

- 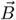 is a vector corresponding to the BOLD time series of a given voxel.
- 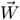 is a vector of regression weights specifying the strength of the link between the voxel and each basis function.
- ***R***_***BF***_ is a matrix containing the response of all BFs to the location-of-interest at any given time.

Each voxel’s RF, then, can be estimated as the linear sum of all basis functions, each weighted by its corresponding regression weight. To this end, we first need to calculate the regression weights vector, W, in equation (1). Thus, we rewrite equation (1) as:

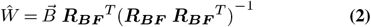

The basis functions are a set of cosine functions centered at uniformly distributed angular positions around a circle, with the radius equivalent to 415 pixels, and raised to the second power. The number (here, 7) and the power (here, 2) in the basis functions formula are chosen to maximize the goodness of fit of the regression in equation (2). Notice that the power is inversely proportional to the width of the basis function. Importantly, due to the fixed eccentricity of the presented stimuli, we configured one-dimensional BFs distribution around a fixed eccentricity instead of two-dimensional BFs covering the whole visual field. This simplifies calculations without any significant effect on the results. We, then, used the same BF set to estimate RFs which result in one-dimensional RFs representing the selectivity profile of each voxel for angular positions at 415 pixel dva eccentricity. Fig. 2A& B show a schematic of the EM and RF estimation. Finally, the RF center and the tuning of each voxel is calculated as the mean (µ) and inverted concentration (*κ*) parameters of a von Mises function fitted to the estimated RF.

### Selectivity maps and regions of interest(ROI)

As not all brain regions exhibit spatial selectivity, we compared spatial tuning between placebo and ketamine conditions exclusively within the spatially-selective areas. We employed a data-driven approach to identify brain regions with significant spatial selectivity. EM calculates the regression weights independently for each voxel. Given that the BFs in the EM are based on well-known spatial selectivity profiles of the visual system, the fit of EM to a voxel’s signal improves as a function of spatial informativeness of the voxel. Therefore, we used EM’s goodness-of-fit (GOF), for each voxel, as a measure of spatial selectivity (but not spatial tuning). Specifically, we measured GOF by the variance explained, which represents the amount of variance in the BOLD signal that can be explained for by the EM.

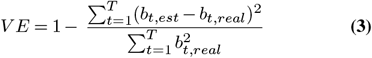

Where b_*t*,*est*_ is the EM-estimated voxel’s bold value at time *t*, b_*t*,*real*_ is the measured voxel’s bold value at time *t*, and T is the number of time points.

To identify spatially-selective ROIs, we first calculated selectivity maps independently for each participant and condition. Next, we averaged these GOF maps across participants and conditions. Then, we z-scored this average GOF map across the whole brain and applied a threshold of 1.65 to create a GOF-based spatial selectivity mask. Finally, we identified spatially-selective ROIs as clusters within this mask that correspond to previously identified retinotopicaly-organized areas across the brain (30, 31) (Fig. 2C).

### Statistical analysis

We compared the EM-estimated tunings under placebo and ketamine administration at the group level. This comparison focused on the ketamine-induced changes in the estimated RF width (*σ*), which is inversely proportional to the concentration parameter (*κ*) of the von Mises function (Fig. 2B). The analysis was conducted by averaging the ketamine-induced change within each ROI and testing against zero. To achieve this, we employed a bootstrapping process with 10,000 permutations to generate a t-distribution of the ketamine-induced changes in *σ*. We then computed the associated p-values to assess the significance of these changes.

To assess the significance of correlation measures, we employed a non-parametric randomization test. Specifically, we compared the intact Pearson correlation coefficient (i.e., the correlation between non-permuted variables) with a corresponding null distribution. This null distribution was generated by calculating correlations between the intact order of one input variable and a permuted version of the second input variable, repeating this process 10,000 times.

To evaluate the significance of the proportion of voxels with positive contrast between placebo and ketamine neural activation, we used a binomial test. Each voxel was considered a sample and a *success* was defined as a voxel exhibiting a positive contrast (i.e., larger activation under placebo, compared to ketamine).

The proportion of success, calculated as the proportion of positive-contrast voxels relative to the total number of voxels in a set, was compared between two voxel sets: i) voxels within each spatially selective ROI, and ii) voxels outside the GOF-based mask (Fig. 3B). The out-of-GOF-mask set was used as the known proportion (p_0_) in the binomial test.

## Data Availability

## Code Availability

All code used in this paper are available from the corresponding author upon reasonable request.

## Supplementary Figures

**Fig. S1.**
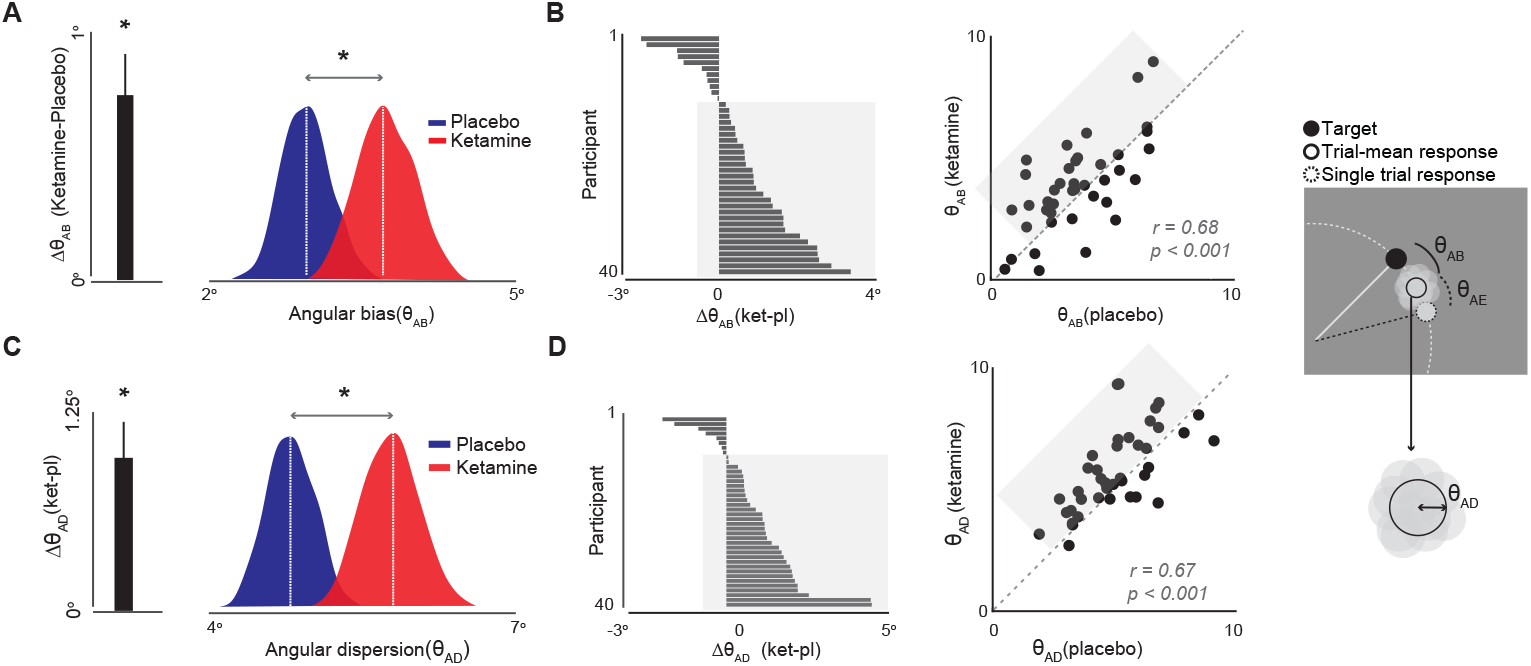
Spatial working memory (sWM) angular bias. **(A)** Spatial working memory (sWM) angular bias; sWM performance was calculated as the combination of two components: angular bias (*θ*_*AB*_) and angular error (*θ*_*AE*_). Angular bias represents the systematic deviation in participants’ responses and is calculated as the average directional distance from the target location across trials(43). Each white circle indicates the response for a single trial. To calculate angular bias, we rotated the target and response of each trial, maintaining their relative distances, so that all trial targets are positioned at the same location (the solid black circle). Angular error is defined as the angular distance between the participant’s response and the target location, adjusted for the angular bias. The bar graph shows the group-mean ketamine-induced change in angular bias, indicating that ketamine significantly increases the sWM angular bias at the group level. The ridgeline plot shows the distribution of angular bias across all participants under placebo and ketamine (dashed white lines indicate group means). **(B)** The ordered histogram shows the worsening effect of ketamine on angular bias across individual participants, indicating that 29 out of 40 participants had worsened angular bias under ketamine compared to placebo (the highlighted area). Each bar represents a single participant, and larger values indicate a stronger effect of ketamine on the angular bias of sWM. The scatter plot depicts the relationship between angular bias under ketamine and placebo conditions. Participants with larger angular bias under placebo have larger error under ketamine, showing that ketamine does not affect the overall distribution of sWM performance across participants. The highlighted rectangle shows participants with worsened angular bias under ketamine, corresponding to the highlighted participants in the ordered histogram. **(C & D)** Spatial working memory (sWM) angular dispersion (plots show similar measures in panels A and B); Another measure of sWM performance, in addition to angular error and angular bias, is sWM dispersion (*θ*_*AD*_). Dispersion is inversely proportional to each participant’s overall sWM precision, reflecting the consistency of sWM responses across repeated attempts. sWM was more dispersed under ketamine, compared to placebo, in 31 out of 40 participants (the highlighted area in the scatter plot in panel D).

**Fig. S2.**
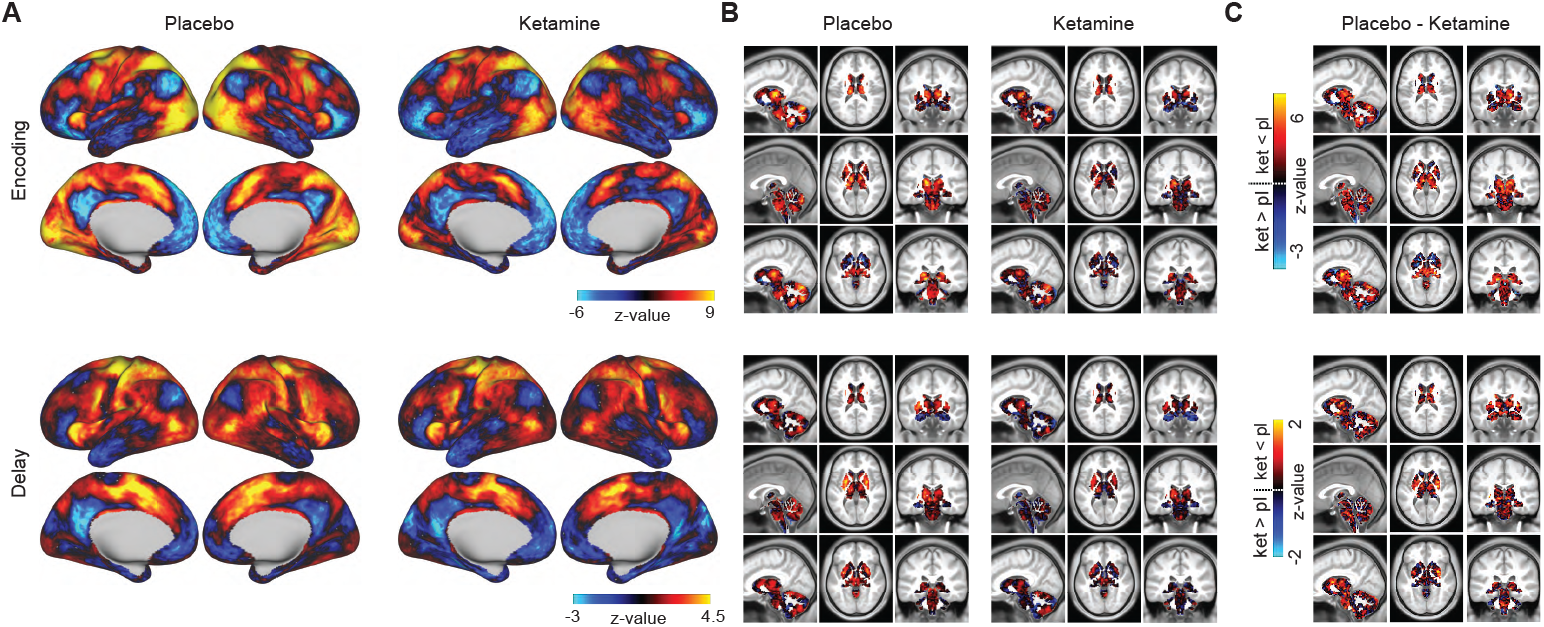
Cortical and subcortical activation. **(A)** Cortical activation during the encoding and delay phases of the task and under placebo and ketamine administration. **(B)** Activation in subcortical regions during the encoding and delay phases of the sWM task under placebo and ketamine conditions (represented through the z-scored t-statistics). **(C)** The ketamine-induced change in encoding- and delay-related activations measured as the difference between activation under placebo and ketamine. The warmer colors on the activation change map indicate stronger deactivating effects of ketamine relative to placebo.

**Fig. S3.**
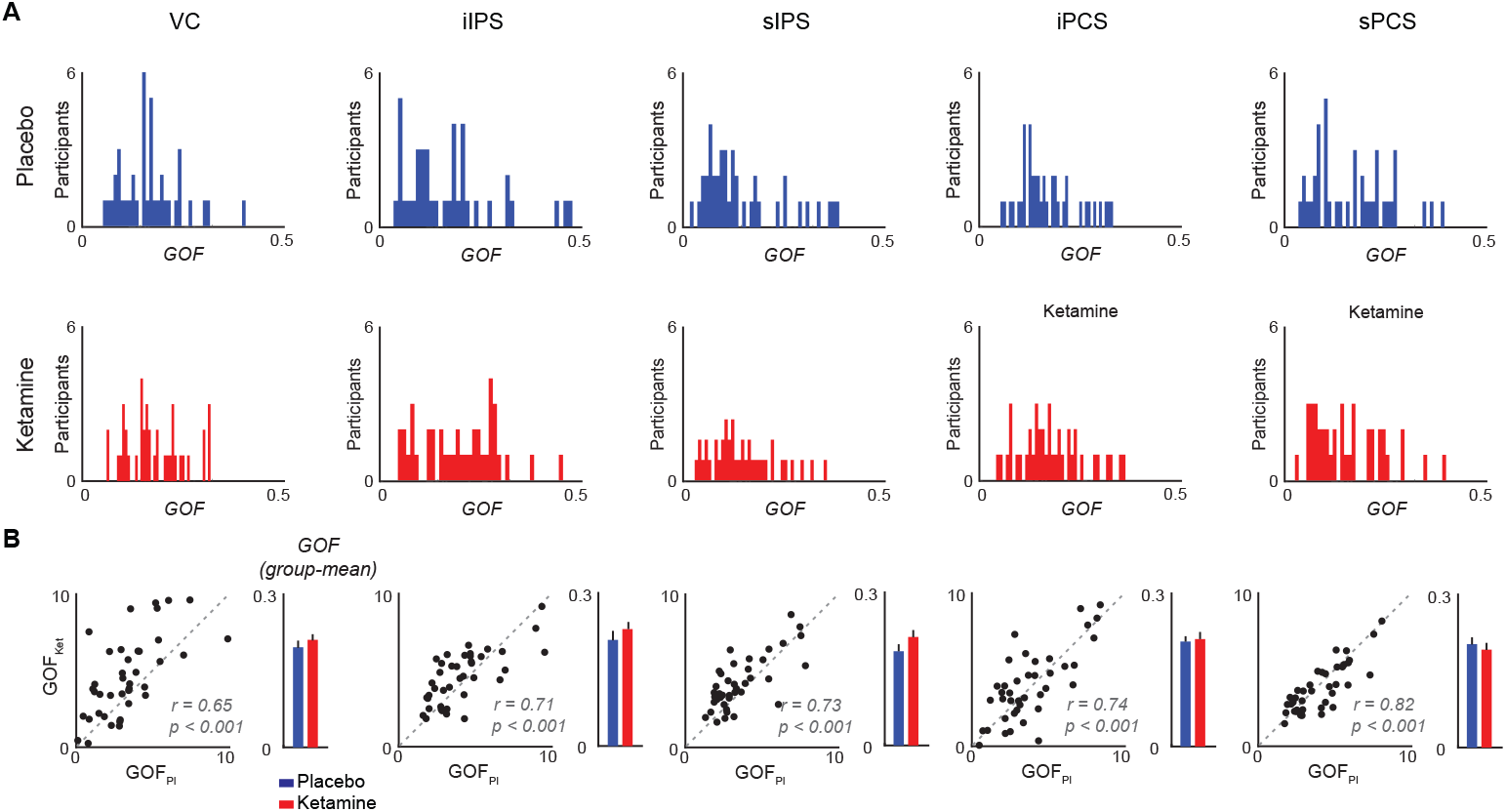
Distribution of goodness-of-fit across participants and reliability of EM estimations. **A)** The upper and lower rows depict the distribution of goodness-of-fit (GOF) of the EM model across individuals under placebo and ketamine administration, respectively, within different spatially-selective ROIs. Each ROI’s GOF is calculated as the average of single voxel GOFs across all voxels within the ROI. **(B)** Scatter plots: the relationship between GOFs under placebo and ketamine within different spatially-selective ROIs. barplots: the group-mean GOF under placebo and ketamine for different ROIs. Error bars show one standard error across participants. Ketamine administration does not change the distribution of GOFs across individuals in any of the ROIs. Overall, ketamine does not significantly affect GOF within the spatially-selective ROIs.

**Fig. S4.**
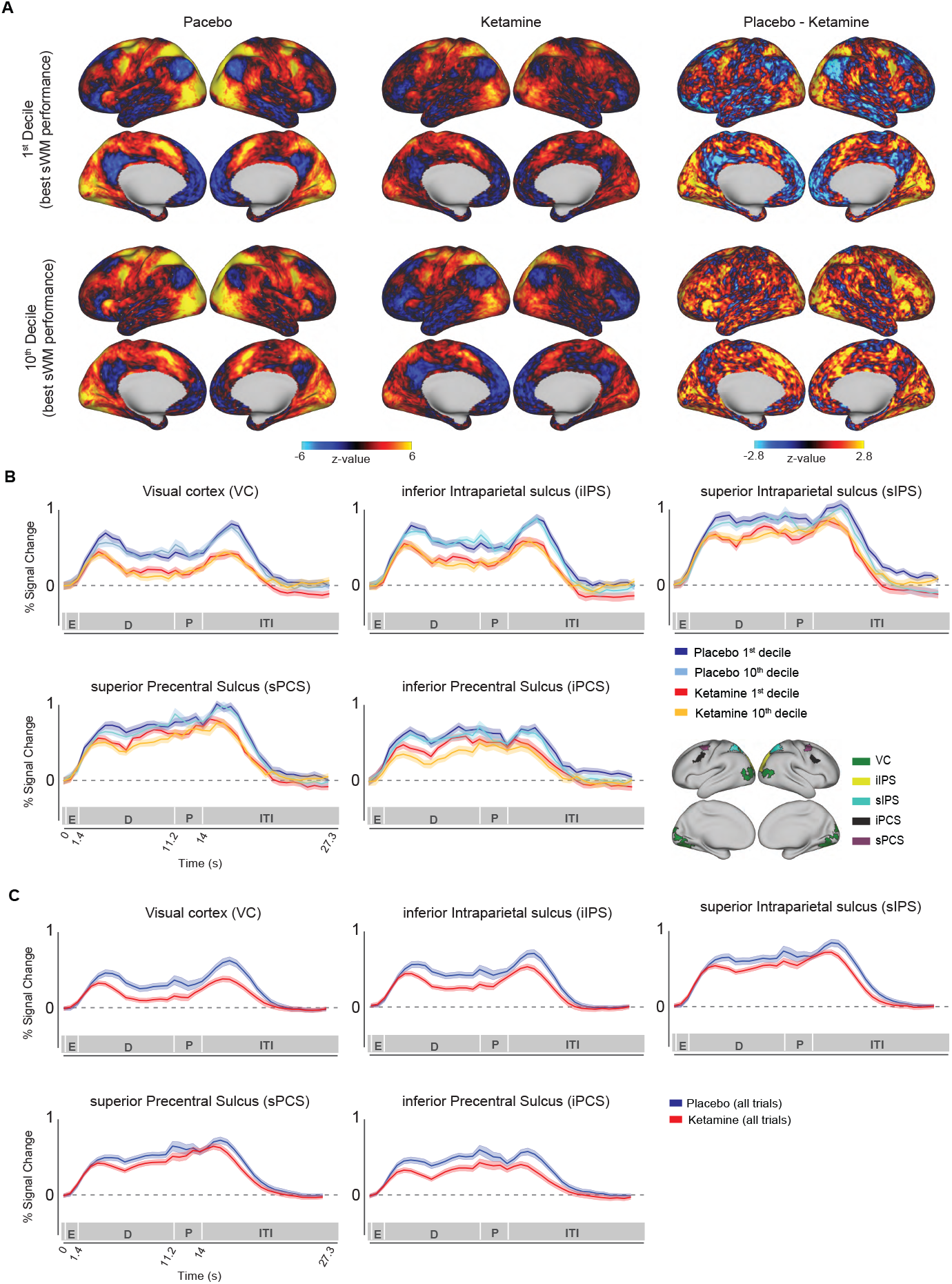
Association between the neural effects of ketamine and its effect on trial-wise sWM accuracy. We tested if neural activation can capture the variability in the trial-wise accuracy of sWM (Fig. 3, main text). **(A)** Neural activation corresponding to placebo and ketamine 1^st^ and 10^th^ deciles (the two left columns) and the ketamine-placebo contrast in activation for each decile. **(B)** Temporal profile of the group-mean activation for the placebo and ketamine 1^st^ and 10^th^ over the course of a trial in all five spatially-selective regions of interest. Shaded areas show one standard error(SE) across participants. **(C)** Temporal profile of the group-mean activation for all trials under placebo and ketamine.

**Fig. S5.**
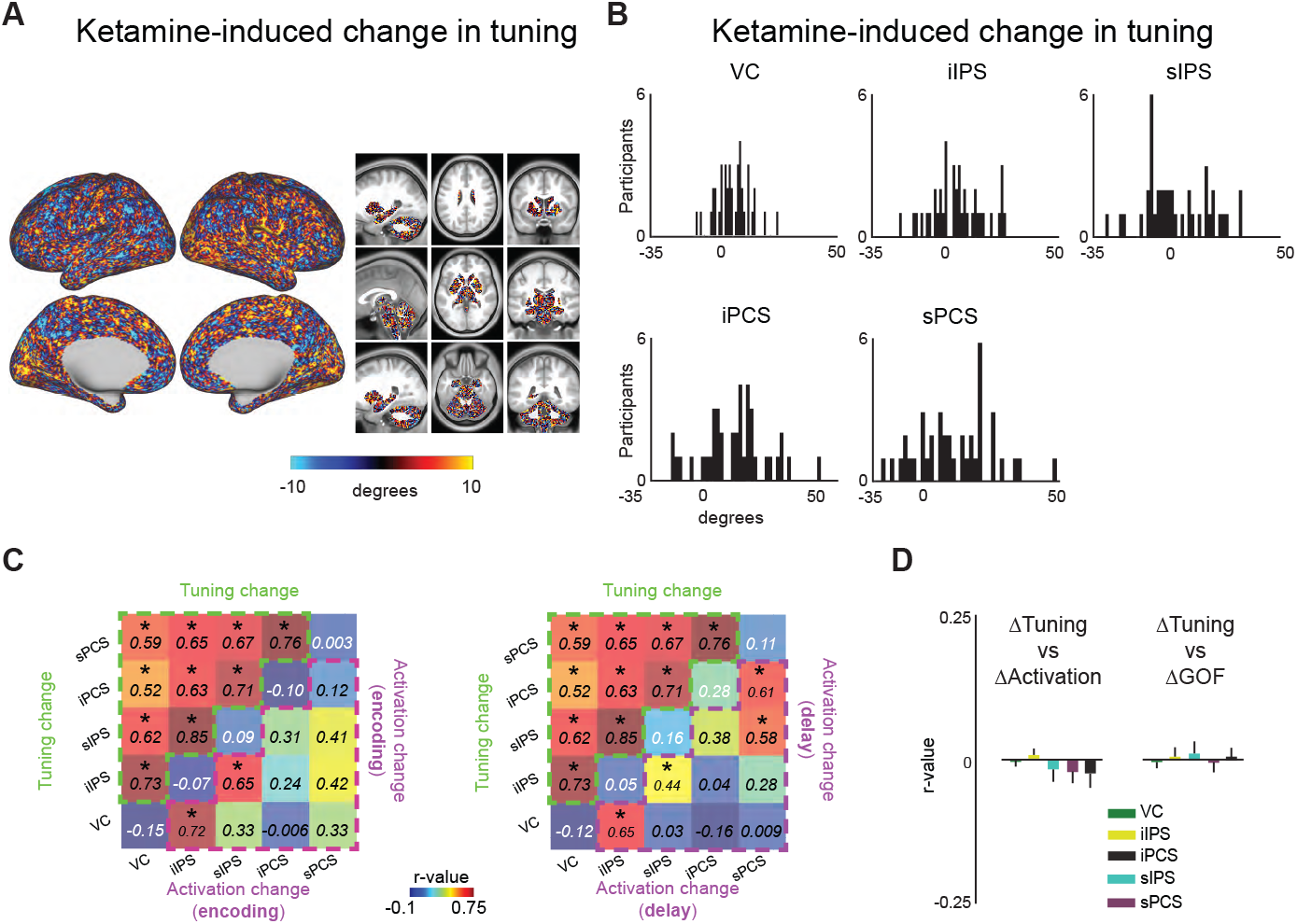
Ketamine-induced change in tuning. **(A)** Ketamine-induced change in tuning across the whole brain, averaged across all participants. **(B)** The distribution of ketamine-induced change in tuning across participants, averaged across all voxels within each spatially-selective ROI. **(C)** The between-ROIs relationship between voxel-average ketamine-induced changes in tuning, the magnitude of activation across ROIs, as well as the within-ROI relationship between tuning and activation; The upper triangle in each matrix contains the across-participants Pearson correlations between ketamine induced changes in tuning for different pairs of spatially-selective ROIs. The lower triangle in each matrix shows the same correlations for ketamine-induced changes in overall neural activation. The diagonal entries represent the across-participants Pearson correlation between ketamine-induced changes in tuning and activation within each ROI. The left and right panels correspond to activation during the encoding phase and the last 4.2 seconds of the delay period, respectively. Asterisks indicate statistically significant correlations (p < 0.05, corrected for multiple comparisons). Note that the upper triangles in both panels are identical, as neural spatial tuning was calculated using only signals from the encoding phase. **(D)** The group-average voxel-wise relationship between ketamine-induced change in tuning and the magnitude of neural activation; Each bar represents the voxel-wise correlation between ketamine-induced change in spatial tuning and ketamine-induced change in the magnitude of BOLD signal (the left set) and change in spatial selectivity (the right set) within an ROI. The height of the bars and error bars represent mean and one standard error across participants, respectively. There is no significant voxel-wise correlation in any of the ROIs.

**Fig. S6.**
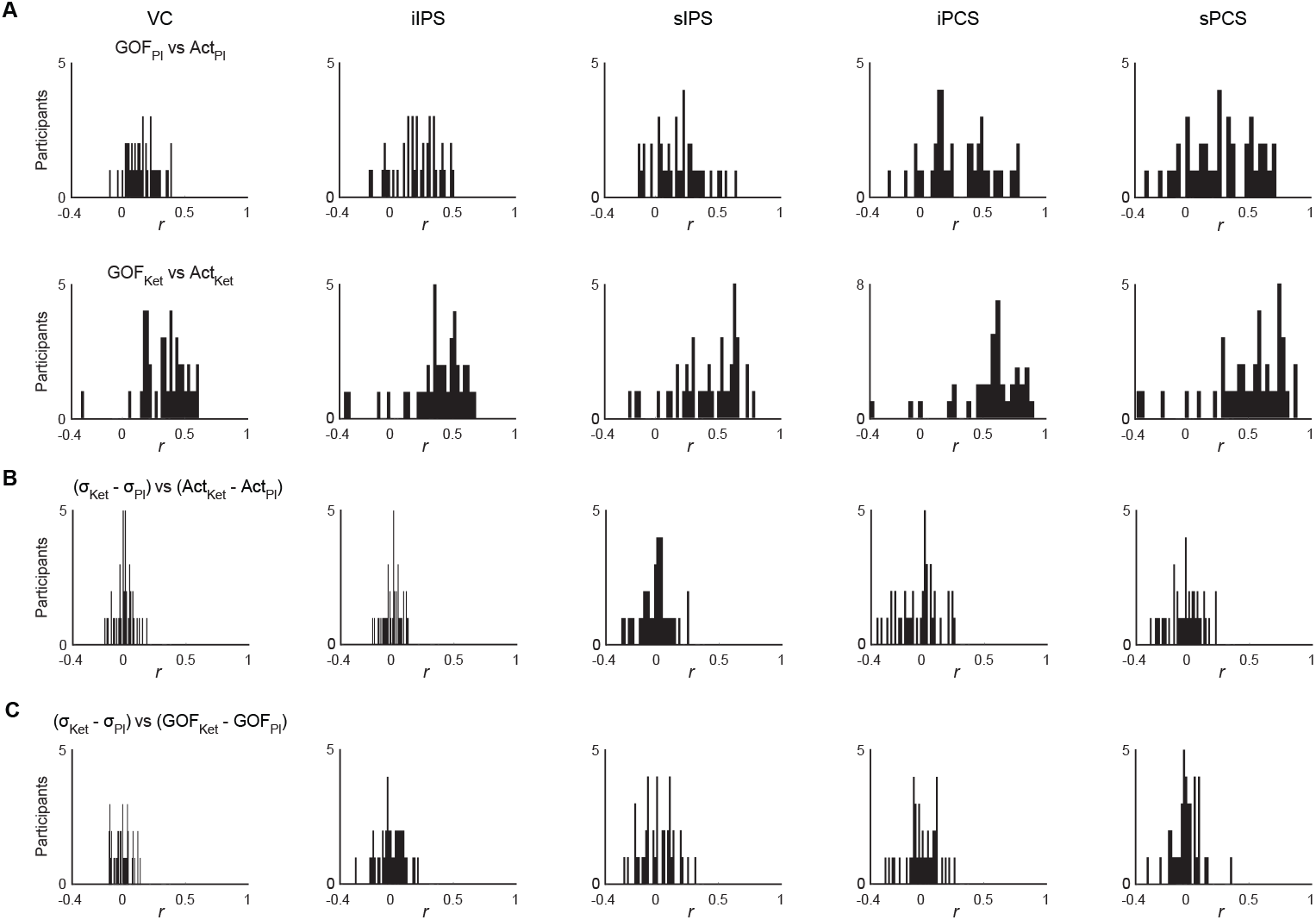
Relationship between neural activation, goodness-of-fit and tuning. **(A)** The distribution of individual-level correlations between goodness-of-fit (GOF) and encoding-related neural activation across participants under placebo (upper row) and ketamine (lower row) within spatially-selective ROIs. For each participant and ROI, the *r-value* is calculated as the correlation between the GOF and activation across all voxels within that ROI. **(B)** The distribution of individual-level correlations between ketamine-induced change in spatial tuning and encoding-related activation within different ROIs. For each participant and ROI, the *r-value* is calculated as the correlation between the ketamine-induced change in spatial tuning and activation across all voxels within that ROI. **(C)** The distribution of individual-level correlations between ketamine-induced change in spatial tuning and GOF within different ROIs. For each participant and ROI, the *r-value* is calculated as the correlation between the ketamine-induced change in spatial tuning and GOF across all voxels within that ROI. Together, panels B and C show that the observed broadening effect of ketamine on spatial tuning is not an intrinsic result of either weaker overall neural signal or the spatial information in the neural signal (measured as the GOF) under ketamine.

